# Quantitative Phase Imaging of Epithelial Monolayer Dynamics

**DOI:** 10.64898/2026.01.17.700037

**Authors:** Silja Borring Låstad, Nigar Abbasova, Thomas Combriat, Dag Kristian Dysthe

## Abstract

This study uses two different quantitative phase imaging techniques (QPI) and for the first time measures the height, volume, and mass dynamics of Madin-Darby Canine Kidney (MDCK) monolayers. We demonstrate novel methods to determine the height of confluent monolayers of cells from 2D and 3D QPI data and validate that the two methods agree. We developed a novel cell segmentation method adapted to QPI images of confluent cell layers and present a robust measure of relative error. We also demonstrate that height statistics of cells can be obtained without segmenting the images. We obtain the following precisions of cell density (1 %), height (3 %), area (5 %) and volume (6 %). Cell height varies 15-25 % over a monolayer and increases 50-100 % when cell density doubles. The average refractive index and the dry mass fraction of the cells, on the other hand, are constant over the entire density range.

## 1. INTRODUCTION

The study of confluent epithelial monolayers in vitro is important to understand the barrier function of epithelia, dynamic phenomena such as wound healing and morphogenesis, and as a model system for active matter. Until now, imaging of epithelial monolayers has been performed with phase contrast or fluorescence microscopy, yielding only 2D information.

The monolayer area and height reflect single-cell volume regulation, where the size parameters (volume, dry mass, and surface) are homeostatically controlled and coupled across timescales. On fast scales (seconds to minutes), mechano-osmotic feedback and pump–leak regulation couple membrane/cortical tension to ion and water flux. On slower scales (hours), metabolic regulation changes impermeable biomass, and volume correspondingly adjusts to preserve intracellular concentrations (dry mass density), coordinating growth with the cell cycle.^1–4^ To date, the proposed volume-regulation pathways have not been systematically compared to data for epithelial monolayers.

Madin-Darby canine kidney (MDCK) cells are canonical examples of cells that form epithelial sheets for the study of collective cell motion.^5–7^ The few existing measurements of cell height and volume of confluent MDCK monolayers by confocal fluorescence microscopy yield conflicting results.^8–13^ This clearly indicates that new methodologies are required to accurately determine the volume-area-height relationships of confluent epithelial monolayers on different timescales.

We report here the use of label-free quantitative phase imaging (QPI) of MDCK epithelial monolayers. We used both 2D QPI and tomographic 3D QPI to study cell volume, cell height, and cell dry mass evolution during the evolution of the MDCK monolayer. The tomographic 3D QPI has access to all these measures at high spatial resolution and produces quantitative measures of fluctuations. Data from the much cheaper and order-of-magnitude faster 2D QPI instrument are compared to tomographic 3D QPI for cross-validation of the results and for future interpretation of 2D QPI data of epithelial sheet dynamics.

## 2. MATERIALS AND METHODS

### 2.1 Quantitative phase imaging

QPI utilises the phase shift of light passing through a cell to estimate its height. Light passing through a cell along the path *s*, gets a phase shift relative to light only passing through water

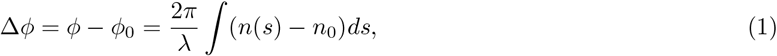

where *λ* is the wavelength of light, *n*_0_ = 1.33 is the refractive index of water, and *n*(*s*) is the refractive index along the path inside the cell. It has been shown that the refractive index is proportional to the volume fraction of non-aqueous (“dry”) matter (proteins, lipids and other biomolecules) *v* in the cell:^14^

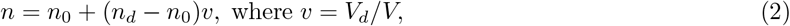

and *n*_*d*_ is the mean refractive index of dry matter, *V*_*d*_ is the volume of dry matter, and *V* is the total volume of the cell.^14^ Since we generally do not know the value of *n*_*d*_ for MDCK, we set it equal to the highest refractive index in our raw data, *n*_*d*_ ≈ 1.43 ± 0.01, which is the refractive index of the lipid vesicles inside the cells.

### 2.2 2D QPI

2D QPI measures the phase shift along the axis perpendicular to the image plane (See Figure 1a). Since 2D QPI does not have separate information about cell height and refractive index, it is customary to use a typical average cell refractive index 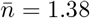 to calculate the 2D QPI height *h*_2*D*_(*x, y*) on the image field (*x, y*):

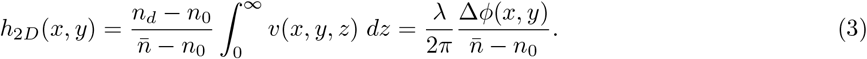

**Figure 1.**
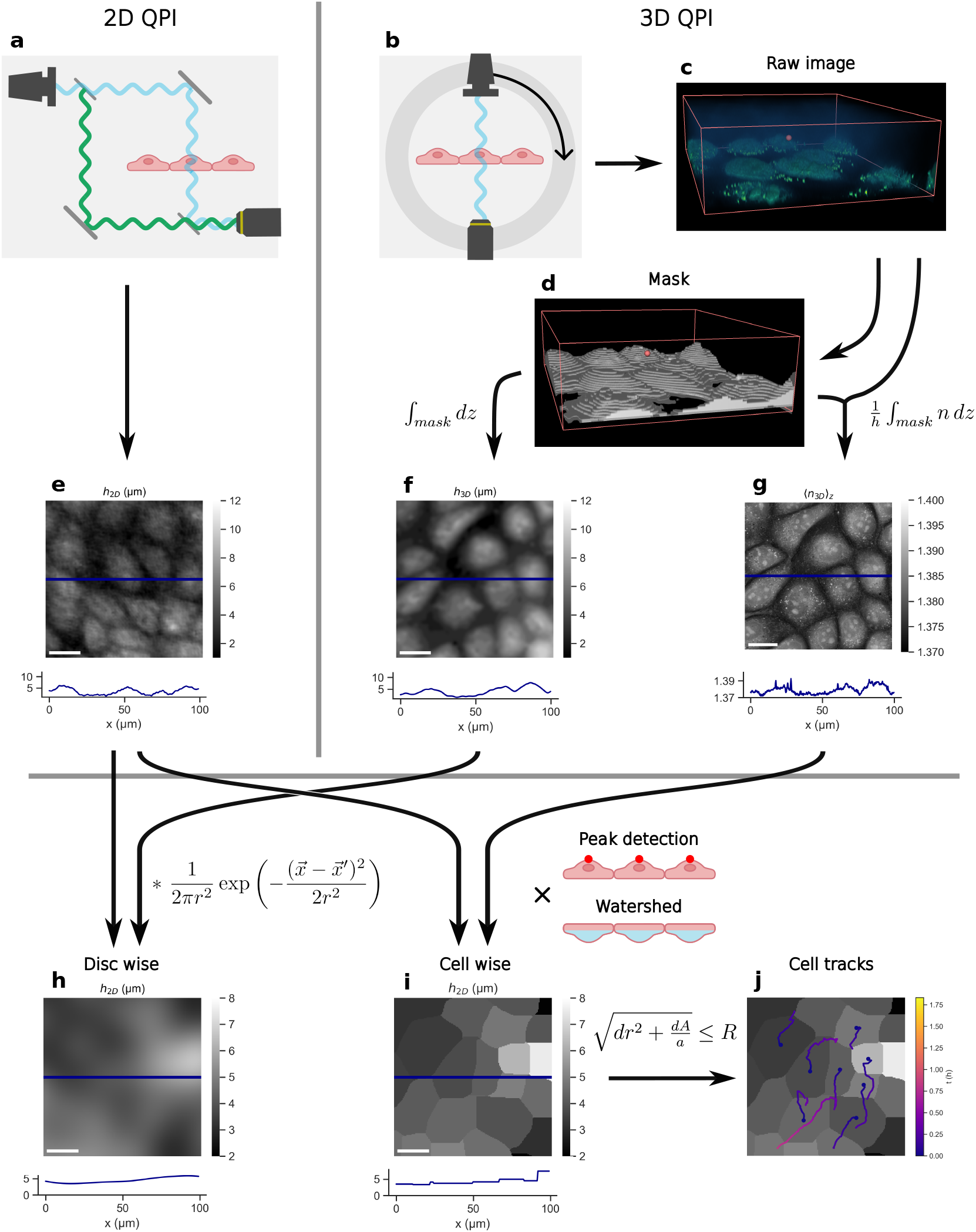
Schematic of image acquisition and preprocessing. **a:** Illustration of 2D QPI. Light passes through the sample and gets a phase shift compared to the reference. **b:** Illustration of 3D QPI. Light passes through the sample from several angles and as such obtain 3D information of the refractive index. **c:** 3D rendering of the raw refractive indices as measured by 3D QPI. **d:** The segmented 3D mask of c. **e:** Cell heights as measured by 2D QPI with the height profile is plotted below. The scalebar represents 20 µm. **f:** Cell heights as measured by 3D QPI. **g:** Height averaged refractive index as measured by 3D QPI. **h:** Disc-wise measures of cell heights are obtained by applying a Gaussian blur with *σ* = *k* · *r* to the heights. **i:** Average cell height of segmented cells are obtained by applying peak detection, then watershed to the heights. **j:** Cell tracks are obtained by minimizing the displacement of the cell centre (*dr*) and the change in cell area (*dA*) scaled by a factor *a*, within a search range *R*.

Integrating *h*_2*D*_ over a whole cell one finds that ∫_*A*_ *h*_2*D*_(*x, y*)*dA* ∝ *V*_*d*_. This indicates that 2D QPI measures the cell dry mass irrespective of the water volume in the cell and cannot distinguish between cells changing projected area while conserving volume or height as long as the dry mass is conserved.

2D QPI measurements were performed with HoloMonitor M3 (Phase Holographic Imaging AB, Sweden, www.phiab.com), and 16 bit unwrapped *h*_2*D*_ images were saved to disc for further treatment by our own Matlab and Python programmes. The time-lapse images from each experiment were registered and minimally cropped to avoid areas without phase information.

#### 2.2.1 Reference phase, *ϕ*_0_

There are two preprocessing steps necessary when using equation (3), phase unwrapping because cos(*ϕ*) = cos(*ϕ*+2*πk*) for all integers *k*, and determining the correct reference phase, *ϕ*_0_ for each image. Phase unwrapping is performed by requiring that there are no phase differences greater than 2*π* between neighbouring pixels and is performed by the HoloMonitor software before presenting *h*_2*D*_ images. The HoloMonitor software calculation of the reference phase assumes distributed cell-free areas to determine the reference phase. We performed 2 different types of experiments that require additional treatment of the reference phase.

**Circular fibronectin patches** of 550 µm diameter confine the MDCK cells to attach and move only inside the field of view. The corners of the images do not have cells and can therefore be used to fit a reference plane *ϕ*_0_(*x, y*).

**Unconfined cell monolayers** on glass covering (more than) the entire field of view requires a different method. We use the histogram *f* (*ϕ*) of unwrapped phases *ϕ* to find the reference phase. We fit a straight line *ŷ* = *aϕ* + *b* to the left flank of the logarithm of the histogram *y* = log *f* (*ϕ*) and set *ϕ*_0_ = − *b/a* to be the intercept with the *x*-axis. This corresponds to shifting the histogram along the *x*-axis for the linear fit to cross the *x*-axis in *x* = 0 (see left hand side of Figure 2).

**Figure 2.**
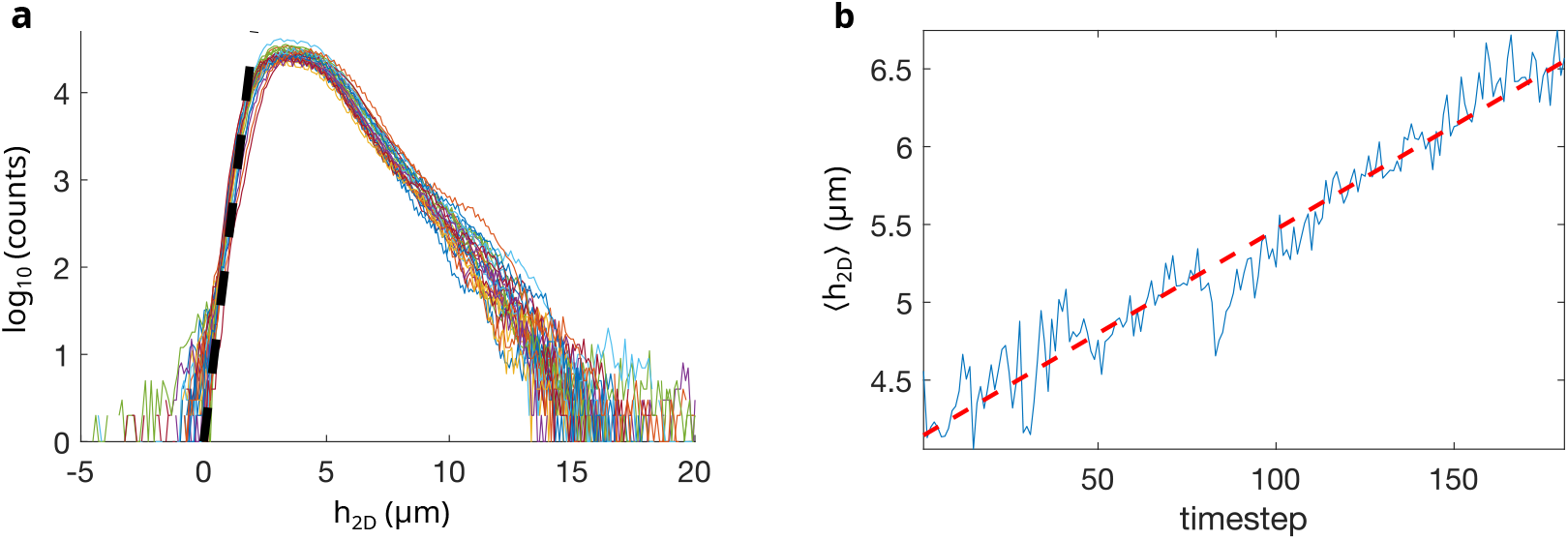
Preprocessing of *h*_2*D*_ . **a:** Zero reference determination. Histograms of pixel heights in 30 images of confluent MDCK II monolayers after first step of reference determination. Black dashed line indicates straight line fit to data on left flank of distribution. The distribution is shifted along *x*-axis for linear fit to cross the *x*-axis in *y* = 0. **b:** Noise removal. Blue line denotes mean monolayer heights ⟨*h*_2*D*_ ⟩ in 181 consecutive images after first step of zero reference determination. Red line denotes linear fit to data.

For time series of 567 × 567 µm^2^ images of the same monolayer, the resulting average height fluctuates around a slowly varying trend with a standard deviation *<* 0.2 µm (see the right hand side of Figure 2). Since there is no reason to believe that the dry volume of cells changes abruptly in time, we adjusted the reference phase to make the average height ⟨*h*_2*D*_⟩ smoothly varying with time (red dashed line on the right hand side of Figure 2). The final pixel heights *h*_2*D*_(*x, y*) have precision ±0.1 µm. The registered and preprocessed height images are saved as 16 bits unsigned integer Tiff files where the pixel values correspond to heights with unit 10 nm.

### 2.3 3D QPI

3D QPI or “tomographic QPI”^15^ measurements were performed with Tomocube HT-X1 (www.tomocube.com) and the reconstructed refractive index volumes *n*(*x, y, z*) were saved to disk for further treatment by our own Python programmes. The Tomographic QPI acquires many images of the phase of light that has passed through all parts of the cell in different directions to reconstruct the refractive index field *n*(*x, y, z*) within the cell (See Figure 1b-c). We obtained the raw data from Tomocube HT-X1 as uncalibrated 3D stacks of refractive indices; thus, they do not represent the actual refractive indices of the cells. The full image is made up of 16 tiles that are stitched together by Tomocube’s proprietary preprocessing software. Since there are regions of *n*(*x, y, z*) = *n*_0_ inside cells and tomographic artefacts of *n*(*x, y, z*) > *n*_0_ in the medium, cell volume cannot be extracted directly from raw data. Therefore, we use a machine learning model that has been trained on Tomocube HT-X1 data^16^ to predict the probability *p*_*cell*_(*x, y, z*), that a voxel belongs to a cell. Note that this model works on the individual 2D slices of a stack and as a consequence 3D information is not part of the input features.

To segment cells from *p*_*cell*_, we first determine the *z*-position of the petri dish, *z*_0_, and then determine *p*_*c*_, the classification threshold for when a voxel is considered to be part of a cell. The estimation of *z*_0_ is illustrated in Figure 3. Figure 3a shows the mean projection of the raw refractive indices along the *y*-axis for a single frame. The real monolayer is located mainly within *z* ∈ [7, 14], but we observe tomographic artefacts above and below this range. *z*_0_ is estimated as the first inflexion point of 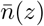, which physically corresponds to the *z*-slice where the image becomes “sharp”. In Figure 3b-c we plot *z* against the mean refractive index along the *z*-axis, 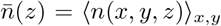, and its derivative, respectively. To account for any tilt of the petri dish, we fit a linear plane to the cell mask at *z*_0_ and subtract this from every tile making up the full image. Since the focus is observed to vary slightly between tiles, the intercept of the plane with each tile is weighted by the fraction of voxels belonging to the cell mask below *z*_0_, such that *z*_0_(*x*_*i*_, *y*_*i*_) = *z*_0_ − *w*_*i*_ · *z*_*plane*_(*x*_*i*_, *y*_*i*_), where *z*_*plane*_ is the fitted plane and takes values ∈ [0, 1], and *w*_*i*_ is a tile-specific weight factor ∈ [0, 2].

**Figure 3.**
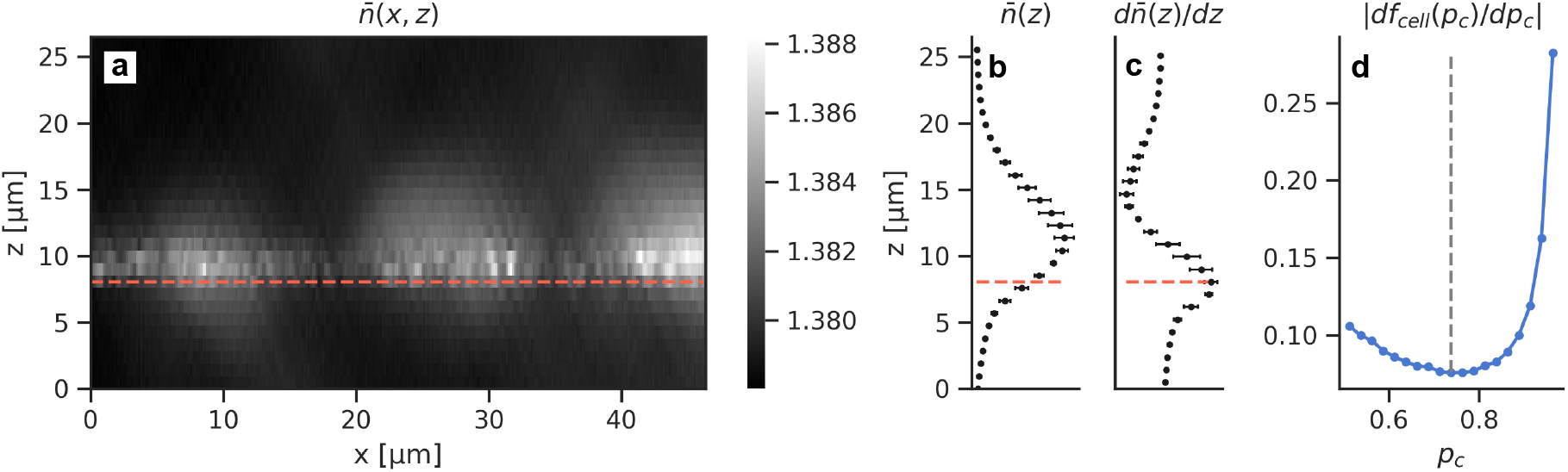
Preprocessing of *h*_3*D*_ . **a-c:** Zero reference determination. **a:** The mean projection of the raw image in the *xz*-plane shows tomographic artefacts above and below the cell layer. The dashed line highlights the zero reference, *z*_0_. **b:** Mean refractive index along the *z*-axis of 40 images. The inflection point of the refractive index is used as *z*_0_. **c:** The derivative of b. **d:** Determination of the classification threshold for the cell mask, *p*_*c*_. The blue dots denote the derivative of the fraction of voxels classified as cells with respect to the classification threshold. The dashed line highlights the minimum, which is chosen as the threshold.

To choose the probability threshold for the machine learning model *p*_*c*_, we consider the fraction of voxels classified as cells *f*_*cell*_(*p*_*c*_), and again choose the inflexion point as the threshold. *f*_*cell*_ is defined as

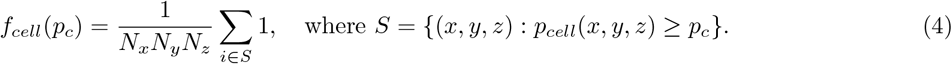

*S* represents the set of all voxels with *p*_*cell*_(*x, y, z*) ≥ *p*_*c*_. The procedure is illustrated in Figure 3d. The classification threshold is specific to each image and typically lies between *p*_*c*_ = 0.55 − 0.75. Finally, we apply a median filter with an ellipsoid-shaped kernel of radii *r*_*z*_ = 3 and *r*_*xy*_ = 11 to the cell mask. An example refractive index field and its cell mask is illustrated in Figure 1c-d. We use this cell mask to obtain *h*_3*D*_(*x, y*) as the extension of the cell mask in *z* for a given position (*x, y*). Then, we smoothen the image with a Gaussian blur with a kernel of 4 × 4 pixels. Similarly to the 2D case, we adjust the average height to be smoothly varying with time. *h*_3*D*_ is, unlike *h*_2*D*_, a direct measure of the physical height of the cell. To confirm that the result is reasonable, we compare *h*_3*D*_(*x, y*) with the number of slices with sharp features in the raw 3D stack for a representative selection of cells. We then use *h*_3*D*_ to measure the height averaged refractive index as:

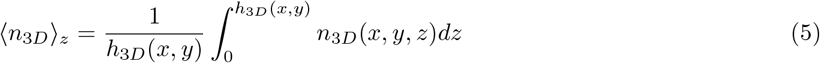

where zero is the base of the cell. Finally, we standardise the uncalibrated distribution of ⟨*n*_3*D*_⟩ _*z*_ by shifting the mean to 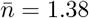, in accordance with the typical cell refractive index used in Eq. (3).

### 2.4 Cell detection and segmentation

The QPI image intensity is proportional to the cell height, which is different from other imaging modalities used for cell imaging. Therefore, standard cell segmentation tools do not work very well for QPI height images of confluent monolayers. Our segmentation method is based on the observation that most cells are highest where the nucleus is and lowest at the cell-cell contact.

We identify individual cells in the monolayer by finding the highest point in each cell. We normalise and smooth *h*_2*D*_(*x, y, t*) (2D QPI) and 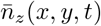 (3D QPI) using a Gaussian blur with standard deviation *σ*_*high*_ and subtract the same image smoothed with *σ*_*low*_ from it to remove large scale variation in the monolayer. The smoothing parameters are collected in Table 1. For *h*_2*D*_(*x, y, t*) we typically use *σ*_*high*_ = 6 pixels (3.3 µm) and *σ*_*low*_ = 12 pixels (6.6 µm) and for 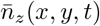 we use *σ*_*high*_ = 35 pixels (5.4 µm) and *σ*_*low*_ = 55 pixels (8.5 µm). Then, we use `scipy`’s peak finder algorithm to detect cells in the smoothed image. We require that the distance between the peaks is larger than *r*(*t*) = *r*_0_ · 2^−*t/*2*τ*^, where *τ* is proportional to the cell doubling time, i.e. the average peak distance decreases inversely proportionally to the cell growth.

**Table 1.**
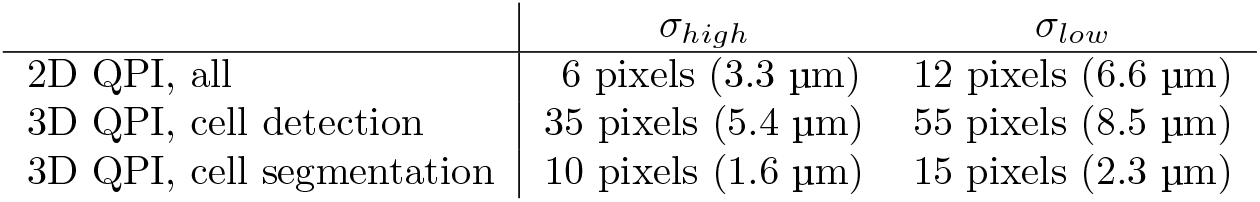
Cell segmentation smoothing parameters.

Since the lowest point between cells typically corresponds to the cell-cell contact, we use a watershed algorithm to delineate which area belongs to which cell, and thereby obtain a mask for every cell in an image frame. The watershed for 3D QPI is performed on 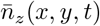 smoothed with *σ*_*high*_ = 10 pixels (1.6 µm) and *σ*_*low*_ = 15 pixels (2.3 µm) and for 2D QPI the smoothing is as for cell detection. The detected cells and their cell areas are illustrated in Figure 1. For each cell, we then compute its total area *A*_*cell*_(*t*), total volume *V*_*cell*_(*t*) = ∑_*x,y*∈*A*_ *h*(*x, y, t*), and mean height 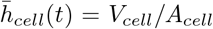. For the tomographic data we also compute the mean refractive index 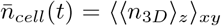. We remove spurious cells with outlier data relative to the area, volume, or height distributions of each dataset.

Finally, we track segmented cells using `trackpy`.^17^ We use both cell position and cell area as tracking parameters so that for every cell, we minimise the distance between points in parameter space

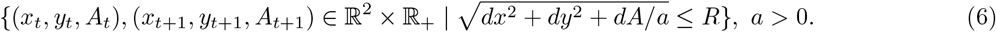

Here, *a* is a rescaling factor that is chosen such that *dA* is in the same range as *dx*^2^, *R* is the upper limit on the search range. Tracking excludes most cell division and severe segmentation errors, but we also require that cell height and cell area differ by less than 50 % between consecutive frames.

### 2.5 Pixel-, cell- and disc-metrics

The cell-wise metric of the property *X, X*_*cell*_, is of biological relevance as it measures the average property *X* of each cell. However, cell-wise metrics are based on segmentation of the images, an error-prone process that determines which part of the image belongs to each cell. Therefore, we also use pixel-wise, *X*(*x, y*) and disc-wise, *X*(*r*; *x, y*) metrics.

#### Cell-wise measure

Cell-wise measurements use segmentation masks for each cell. Cell-wise distributions of a property *x* first perform an average of *x* over each cell mask 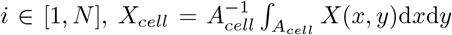 d*x*d*y* and then calculate the distribution, mean, and standard deviation over the *N* cells.

#### Disc-wise measure

Because segmentation is inherently error-prone, we also perform disc-wise measures. Instead of averaging over each cell mask, we first smooth the image with a two-dimensional Gaussian disc with a standard deviation of *r* = *k* · *r*_*cell*_ surrounding each pixel, *X*(*r*; *x, y*) = (*X* ∗ *G*_*r*_) (*x, y*), where 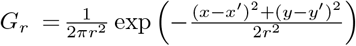 . We then calculate the distribution, mean, and standard deviation over all pixels. Here, *r*_*cell*_ is the average cell radius in a given frame and *k* is chosen such that the disc-wise height variations are similar to their cell-wise counterparts. In this study, we use *k*_2*D*_ = 0.75 and *k*_3*D*_ = 0.95.

### 2.6 Cell culturing

Madin–Darby canine kidney cells, MDCK parental line (ATCC NBL-2 CCL-34, purchased in 2004) were cultured in complete Dulbecco’s Modified Eagle Medium (DMEM, Sigma-Aldrich). All cells were grown in the presence of 10 % FBS (Sigma-Aldrich) and 1 % penicillin/streptomycin (Sigma-Aldrich) at 37 °C in a humidified atmosphere containing 5 % CO_2_. The experiments were carried out at passaging numbers 2-20. Cells were seeded at different densities in 35 mm glass bottom dishes (MatTek, part no: P35G-1.5-14-C) 24-72 hours before imaging was started.

## 3 RESULTS

### 3.1 Cell detection, segmentation, and cell area precision

The uncertainty of cell detection is quantified in terms of the standard deviation of the residuals of a low-pass filter applied to cell density 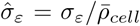. We measure the cell density as the number of detected cells in the field of view (FOV), divided by their total cell area. The cut-off frequency, 1 h^−1^, is then chosen as the lowest frequency that produces uncorrelated residuals. The time evolution of the cell density of a data set is shown in Figure 4a with the residuals plotted below. For cell densities in the range 1600 − 2600 cells/mm^2^, we see that the error in cell count lies within 25 cells/mm^2^. Our cell detection is consistent within an uncertainty of 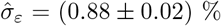, however, it will often fail at detecting cells in very dense regions, and these will often be excluded from the analyses.

**Figure 4.**
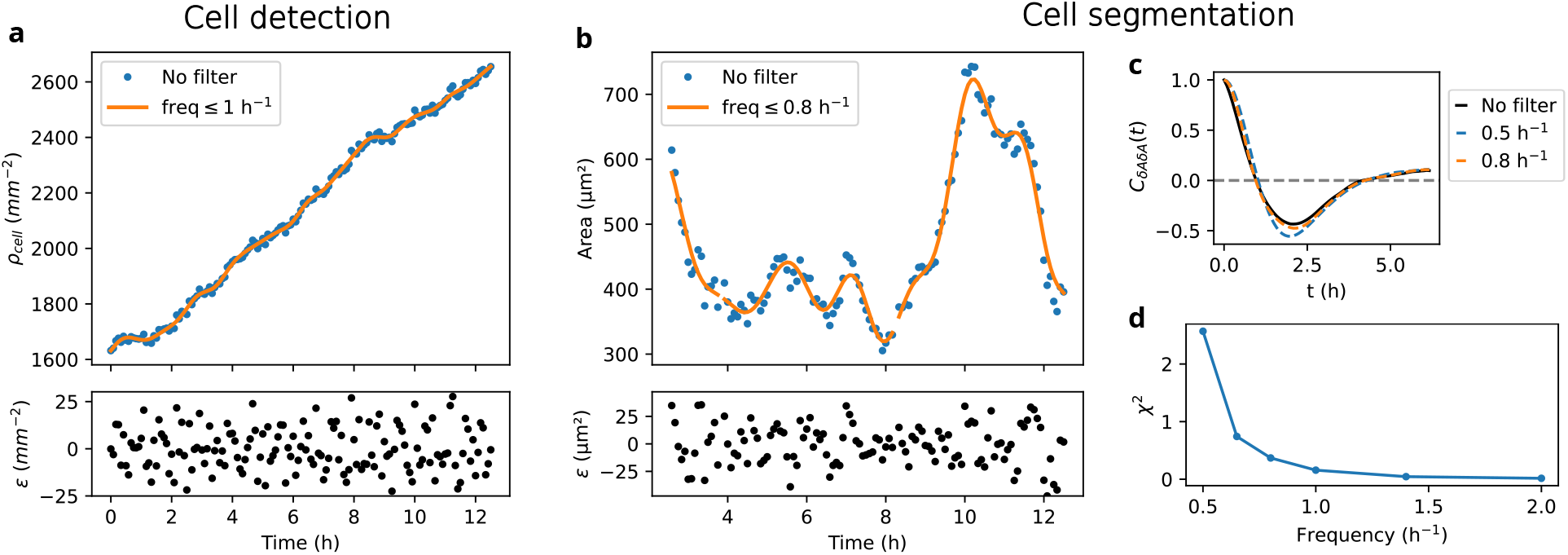
Cell segmentation error analysis. **a:** Time evolution of cell density, *ρ*_*cell*_, before (blue dots) and after (orange line) applying a low-pass filter with cutoff frequency 1 *h*^−1^. The residuals of the low-pass filter are plotted below (black dots). **b:** The cell area of a single cell (blue dots) with the low-pass filtered area plotted on top (orange line). The residuals are plotted below. **c:** The autocorrelation of detrended area fluctuations, *C*_*δAδA*_(*t*)) for the original data (black line), and low-pass filtered data with cutoff frequency 0.5 (blue dashed line), and 0.8 *h*^*−*1^ (orange dashed line). **d:** The effect of the low-pass filtering is quantified in terms of the *χ*^2^ of the filtered and unfiltered data as a function of cutoff frequency).

We perform all analyses without manually correcting segmentations to avoid bias. Although cell area changes on shorter timescales than cell growth, we consider high frequency area fluctuations to be segmentation errors, and therefore apply a low-pass filter to all cell area tracks *A*_*i*_(*t*). The area of a cell with and without the low-pass filter is shown in Figure 4b. The area fluctuates around 500 µm^2^, displaying both small fluctuations with a period of ∼2 h and large fluctuations with periods ≥4 h. Because noise is significantly higher in cell area (Figure 4b), compared to cell density (Figure 4a), we want to ensure that we are only removing uncorrelated fluctuations. The average cell area decreases with cell density; therefore, we fit a straight line to the data *Â*(*t*) and calculate the fluctuations in the detrended area *δA* = *A*(*t*) − *Â*(*t*). We then calculate the time autocorrelation of the detrended area, *C*_*δAδA*_(*t*) averaging over all tracks. We repeat this procedure with low-pass filtered tracks of varying cutoff frequencies. The autocorrelation of two such tracks is plotted on top of the autocorrelation of the unfiltered *δA* (black line) in Figure 4c. For a cutoff frequency of 0.8 h^−1^, the autocorrelation mostly follows the unfiltered data and importantly crosses 0 at equal times as the unfiltered data. With a cutoff frequency of 0.5 h^−1^, we observe that the autocorrelation start deviating notably from the unfiltered autocorrelation. We compute the *χ*^2^ between the filtered and the unfiltered *C*_*δAδA*_(*t*) for all cutoff frequencies. This is shown in Figure 4d, where we see that it increases exponentially as we decrease the cutoff frequency. We use this result to choose a cutoff frequency of 0.8 h^−1^ to obtain the “true area evolution”, *A*^′^(*t*), of each cell, considering the residuals as momentary random errors of cell segmentation. Averaging the standard deviation of the residuals over all datasets, we obtain the precision of *A*^′^(*t*) (4.8 ± 0.2) %.

### 3.2 Validation of height measurements

#### 3.2.1 Absolute heights

The properties of MDCK monolayers are very dependent on cell density, *ρ*_*cell*_.^18, 19^ In Figure 5 one observes that the cell density increases approximately linearly with time. The cell doubling time varies from 7 to 30 hours with a mean of 17 hours. The monolayer properties at a given cell density also depend on the seeding density^20, 21^ and the average cell height and cell doubling time can even vary from region to region within the same dish (see, for example, datasets A1-13 and A1-18 in Figure 5). The average cell volumes ⟨*V* (*t*)⟩ _*cell*_ = ⟨*h*(*t*) ⟩ _*cell*_*/ρ*_*cell*_ tend to decrease slightly with increasing cell density. The black dashed lines correspond to constant volume and thus *h*(*ρ*) = *V/A*(*ρ*) ∝ *ρ*.

**Figure 5.**
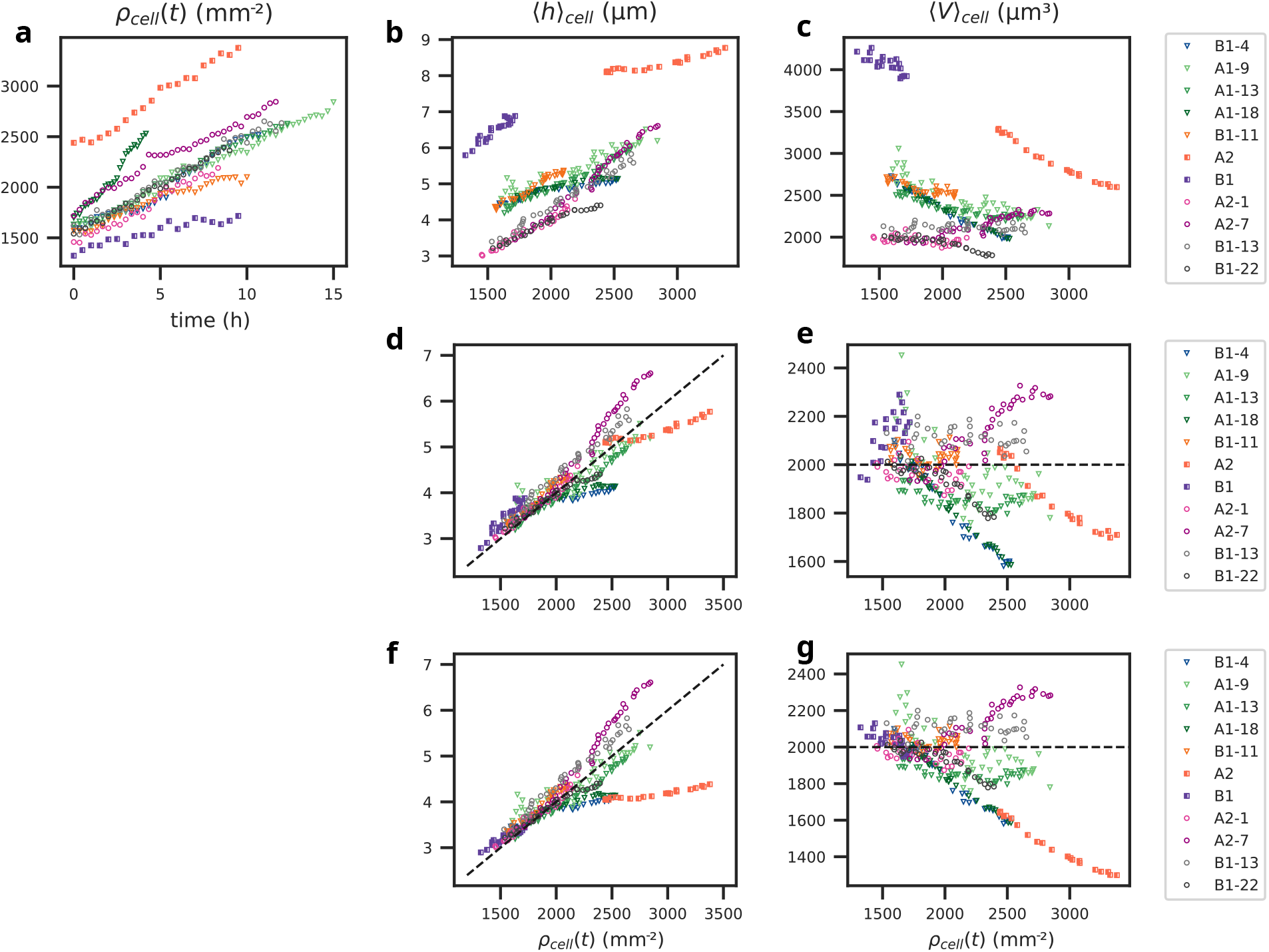
Average density, height and cell volume MDCK monolayers. Squares: 3D QPI on glass, Circles: 2D QPI on glass, Triangles: 2D QPI on 500 µm diameter fibronectin patches. **a:** Cell density evolution in time. **Middle column:** average cell height as function of cell density. **Right column:** Average cell volume (⟨*V* ⟩_*cell*_ = ⟨*h*_*cell*_*A*_*cell*_⟩) as function of cell density. **a-c:** Data treated as described in the methods section. Two bottom rows with adjusted parameters in equation(7) **d-e:** 3D QPI data shifted by *δ*_3*D*_ = −3 µm and 2D QPI data on glass shifted by *δ*_*g*_ = −1 µm. **f-g:** 3D QPI scaled by *γ*_3*D*_ = *γ*_3*D*,0_*/*2 and 2D QPI data on glass shifted by *δ*_*g*_ = −1 µm. *γ*_2*D*_ = *γ*_2*D*,0_.

The upper row of Figure 5 shows the data treated as described in the methods section. Although we know there can be variations due to the initial seeding of cells, it is curious that there are three distinct groups of height and volume depending on the measurement and preprocessing methods (3D, 2D with and without reference area). In first order, we can assume that the measured height *h*_*m*_ is a linear function of the real cell height *h*. The 3D QPI data and the 2D QPI data for cells on a fibronectin patch have a physical zero height reference and therefore we get the following relations between the real and measured heights:

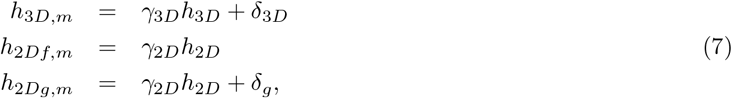

where the subscripts *f* and *g* signify fibronectin and glass. The proportionality factor *γ*_3*D*_ should be 1, but since we do not know exactly how the Tomocube reconstruction algorithm works, we relax this requirement. *γ*_2*D*_ depends on the average refractive index of the cells that we have fixed at 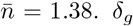 is the absolute error of the zero reference as estimated in Section 2.2. *δ*_3*D*_ is the *z*-resolution of the 3D stack, which here is *δ*_3*D*_ ≤ 0.8 µm. Figure 6 shows the average refractive index as a function of density and shows that it does not change more than 2 % over the entire density range. This is a smaller change than the accuracy of *h*_2*D*_ and we conclude that *γ*_2*D*_ is effectively constant throughout the density range. Since each group of data in Figure 5b seems to show consistent heights as a function of density, we can choose to adjust *γ*_3*D*_/*γ*_2*D*_ by a factor 2 and set *δ*_*g*_ = −1 µm to obtain agreement between the three groups. In order to find the absolute value of the average height of the monolayer, one would have to use a cell phantom of similar and known height and refractive index.

**Figure 6.**
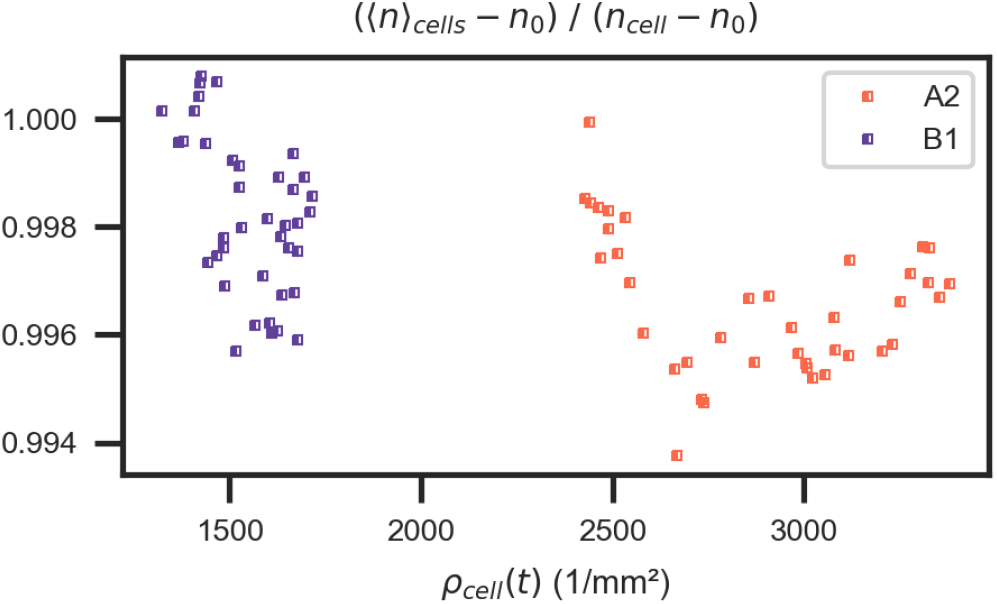
Population average refractive index as function of cell density for two datasets.

Applying these adjustments to the data we observe (see the bottom row of Figure 5) good agreement between all data sets at low density. At high densities, the cell heights on the fibronectin patches are larger than those of the unconfined monolayers on glass, and there is an increasing spread in heights for the unconfined monolayers on glass. In the middle row, we have shifted both 3D and 2D QPI data to obtain the best agreement between the three datasets, although we have no theoretical justification for shifting the 3D data set to as much as *δ*_3*D*_ = 3 µm instead of scaling *γ*_3*D*_.

#### 3.2.2 Relative height variations

Since it is difficult to determine the absolute heights and volumes of cells, we choose to compare relative height variations using 3D and 2D QPI techniques. In Figure 7 we plot the relative height variations over a monolayer, quantified by the relative standard deviation of the heights *σ*_*h*_/ ⟨*h*⟩ _*xy*_. We observe that the variation decreases with cell density, and that the variations measured by 2D and 3D QPI agree very well. We also observe that the disc-wise measurements of local heights (7b) agree very well with the cell-wise measurements(7d). This means that one does not need to rely on segmentation to obtain dynamic information about monolayer height variations.

**Figure 7.**
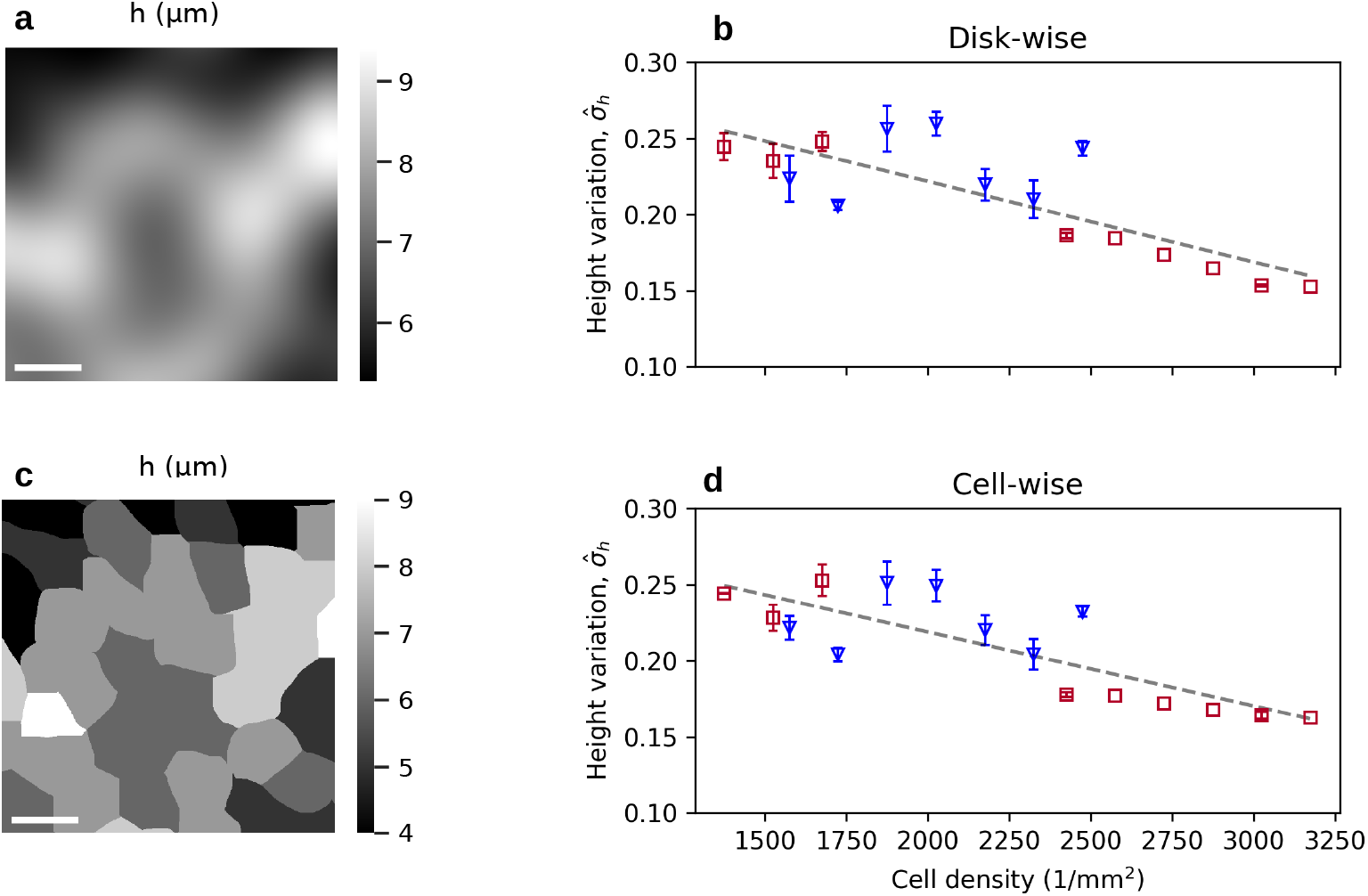
Height variations over a monolayer. **Left:** Illustration of disc-wise heights (a) and cell-wise (c) heights of the same FOV. The scalebars represent 20 µm. **Right:** Spatial height variation in a monolayer as function of cell density, *ρ*_*cell*_ as measured by the disc-wise (b) and cell-wise (d) metric. Red squares denote 3D QPI and blue triangles denote 2D QPI. The dashed line is a linear fit to the data.

## 4. DISCUSSION AND CONCLUSIONS

We have developed a range of methods for post-processing data from the commercial microscopes Holomonitor and Tomocube, to obtain quantitative height, area, and volume data of confluent MDCK monolayers. The absolute heights that we present still have an unknown offset from real heights that can only be determined by calibrating measurements using cell phantoms with known height and refractive index. However, height variations in space and time are robust and quantitative with a measurement precision of 3 % or less. The deviations from the regression line in Figure 7 up to 20 % are due to variations between biological replicates.

We have also developed robust cell detection and segmentation algorithms for QPI data of confluent cell monolayers and evaluated the precision of cell detection and individual cell area determination to be 1 % and 5 %, respectively. Combining height and area precisions, we find that our determination of single cell volumes has a precision of 6 %. We note that there is significant fine tuning to our datasets of the image processing to achieve these precisions. In other datasets, it will probably be useful to use the disc-wise metrics we have presented to avoid the errors introduced by the cell identification and segmentation.

Zehnder et al.,^8, 9^ Thiagarajan et al.^11, 12^ and Devany et al.^13^ have used fluorescence microscopy to measure MDCK monolayer heights. The two first focused on spatial and temporal variations in cell height and volume, the latter focused on long term changes of height and volume with cell density. Thiagarajan et al.^11, 12^ found that the height of the monolayer varies in space and time, whereas other studies found that the height is constant in both space and time. Although we show that the absolute values of the heights measured by QPI can be debated, there is no doubt about the trends:

- Cell height varies significantly (15-25 % depending on density) over a monolayer at any given time (see Figure 7).
- The average height of the monolayer increases 50-100 % when the cell density doubles (see the middle column of Figure 5).
- The average refractive index of the population 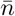 and therefore the average volume fraction of dry mass 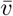 change by no more than 2 % over the entire density range of 1400-3500 cells/mm^2^.

The average cell volume seems to decrease slightly with cell density (see the right column of Figure 5), but there seem to be different trends depending on the substrate being glass or fibronectin. To calibrate the measurements even better than now, one will need to perform unconfined measurements on fibronectin and use cell phantoms with known height and refractive index.

## Acknowledgements

This project has received funding from the European Union’s Horizon 2020 research and innovation programme under the Marie Sklodowska-Curie grant agreement No 945371. The authors thank UiO:LifeScience for the use of the HoloMonitor and the Hybrid Technology Hub, UiO for the use of the TomoCube and Edna (Xian) Hu for providing the MDCK cells.

## Declaration of generative AI and AI-assisted technologies in the writing process

During the preparation of this work the authors used gpt.uio.no in order to improve readability and language. After using this tool, the authors reviewed and edited the content as needed and take full responsibility for the content of the publication.

